# Non-peroxide antibacterial activity of *Meliponula* (*Axestotrigona*) *ferruginea* honey from Tanzania

**DOI:** 10.1101/2024.03.20.585900

**Authors:** Christopher Alphonce Mduda

## Abstract

Honey serves as a medicinal food that is utilized for both preventative care and the treatment of various ailments. Amidst the contemporary challenge of antibiotic resistance, honey emerges as a promising natural antimicrobial solution. The efficacy of honey in therapy hinges on its mechanisms of antimicrobial action. Thus, this study investigated the non-peroxide antibacterial properties of honey sources from a stingless bee species, *Meliponula* (*Axestotrigona*) *ferruginea*, that is commonly managed in Tanzania. The findings reveal that honey from stingless bees exhibits remarkable antibacterial efficacy against both resistant and susceptible bacterial strains. Notably, the studied honey samples retained a substantial portion of their antibacterial potency (89.9 - 98.7%) even after the removal of hydrogen peroxide. Interestingly, the antibacterial activity of honey did not correlate with its total phenolic and flavonoid content, suggesting the influence of specific bioactive compounds rather than overall phytochemical content. Stingless bee honey was most effective against gram-positive bacterial strains, particularly *Staphylococcus aureus*. These results underscore the therapeutic potential of stingless bee honey for the management of pathogenic bacteria. Future investigations should focus on elucidating the specific bioactive compounds present in stingless bee honey to bolster its clinical applications.

## 1.0 Introduction

Bacterial pathogens constantly evolve, acquiring resistance to existing antibiotics (Christaki et al. 2020). This evolutionary arms race complicates treatment strategies, as once-effective antibiotics lose their efficacy against resistant strains (Jalalifar et al. 2024). The overuse and misuse of antibiotics in both clinical and agricultural settings exacerbate this problem, fostering the emergence and spread of drug-resistant bacteria (Endale et al. 2023). Resistant bacterial strains such as Methicillin Resistant *Staphylococcus aureus* (MRSA) have become cosmopolitan due to their capacity to spread rapidly (Nishio et al. 2015). Furthermore, the development of new antibiotics lags behind the emergence of resistance, with few novel drugs reaching the market in recent decades. The discovery of new antibiotics has slowed down over the past years due to high costs of drug research, resulting into few effective antimicrobials which are often associated with high costs and multiple side effects for patients (Cardozo et al. 2013, Zainol et al. 2013). For this reason, the search for novel antimicrobial compounds sourced from natural products is paramount.

Honey is a sweet substance produced by bees from the nectar collected from plants. It contains sugars as the major component, alongside water, organic acids, enzymes, vitamins, proteins, amino acids, minerals, polyphenols and volatile compounds (da Silva et al. 2016). There is a broad array of honey varieties with dinstinct flavor, color, and odor, originating from various floral sources and bee species. Honey has been used as a food and an integral part of traditional medicine since ancient times (Kuropatnicki et al. 2018). The medicinal uses of honey persisted to the modern era, giving rise to an alternative medicine discipline known as Apitherapy, which utilizes honey and other bee products for health treatment (Mandal and Mandal 2011). The medicinal potential of honey is acknowledged for its antimicrobial, antioxidant, anti-inflammatory, anti-cancer, antidiabetic and immunomodulatory properties (Meo et al. 2017). Due its local availability, affordability, and minimal risks of toxicity and microbial resistance, honey stands out as a valuable alternative for treating bacterial pathogens (Mduda et al. 2023e). Currently, several types of honey are marketed as medical-grade honeys with standardized levels of antibacterial activity. The best known is the Manuka honey which is produced from *Leptospermum* species and is reported to be effective against more than 60 species of bacteria (Mandal and Mandal 2011, Nolan 2020).

The diverse components and physical properties of honey collectively contribute to its antimicrobial potency. In an undiluted state, the antimicrobial activity of honey is largely attributed to its high osmolarity and low pH (Zainol et al. 2013). High sugar concentration in honey exerts osmotic pressure on bacterial cells, leading to dehydration and cell shrinkage (Albaridi 2019). Additionally, pH of honey (3.2 – 4.5) is far below the optimal pH for the growth of most bacteria which ranges from 6.5 to 7.5 (Almasaudi 2021). Dilution of honey activate the enzyme glucose oxidase which catalyzes the conversion of glucose to gluconic acid and hydrogen peroxide (Zainol et al. 2013). Hydrogen peroxide is a strong disinfectant which contributes to the antimicrobial efficacy of honey. The maximum level of hydrogen peroxide is achieved when honey is diluted by 30 to 50% (Almasaudi 2021). However, hydrogen peroxide is susceptible to degradation by catalase enzyme in living tissues making it less effective during therapy (Ewnetu et al. 2013, Mduda et al. 2023e). The antibacterial activity of honey can decrease by up to 100-fold following the removal of hydrogen peroxide (Mandal and Mandal 2011). Nonetheless, certain varieties of honey possess non-peroxide activity which allows them to maintain antibacterial potency even after the removal of hydrogen peroxide. The non-peroxide activity results from various elements found in honey, such as phenolic compounds, flavonoids, antibacterial peptides, methylglyoxal, methyl syringate, and other trace components (Zainol et al. 2013).

The use of honey in traditional medicine is widespread in eastern Africa (Kiprono et al. 2022, Mduda et al. 2023d, Héger et al. 2023). To date, various studies have been conducted to investigate the antimicrobial properties of honeys from this region (Ewnetu et al. 2023, Mokaya et al. 2020, Mduda et al. 2023e, Rikohe et al. 2023, Mduda et al. 2024). Findings from Ethiopia and Tanzania revealed that stingless bee honey was more effective against both gram-positive and gram-negative bacteria in comparison to *Apis mellifera* honey (Ewnetu et al. 2013, Mduda et al. 2024). However, little is still known about the mechanisms that underlie the antimicrobial potency of stingless bee honey. The current study investigated for the first time the non-peroxide antibacterial activity of honey samples produced by *Meliponula* (*Axestotrigona*) *ferruginea,* a commonly managed stingless bee species in Tanzania (Mduda et al. 2023d, Mduda et al. 2023b). Specifically, honey samples from Siha and Kibiti districts were tested against resistant and susceptible strains of common pathogenic bacteria. The findings of this study will offer valuable insights into the effectiveness of stingless bee honey as a powerful antimicrobial agent against prevalent pathogenic bacteria, potentially expanding its utility in clinical therapy.

## 2.0 Materials and Methods

### 2.1 Honey samples

Honey samples were gathered from *Meliponula* (*Axestotrigona*) *ferruginea* hives from two districts, namely Siha and Kibiti. Siha district, situated in the northern highlands of Tanzania, featured sampling locations at the western foothills of the Mount Kilimanjaro. This area boasts Afromontane vegetation, known for its multi-layered, evergreen flora, spanning elevations between 1,200 to 2,500 meters with remarkable plant diversity (Foley et al. 2014). Conversely, Kibiti district lies along the eastern coast region, where honey samples were collected from the Rufiji Delta. This delta is renowned for hosting the largest concentration of mangroves on the eastern coast of Africa, representing six distinct families: Avicenniaceae, Combretaceae, Meliaceae, Rhizophoraceae, Sonneratiaceae, and Sterculiaceae (Monga et al., 2018). At both locations, stingless bee hives were managed within semi-natural settings, characterized by the natural vegetation. Sample collection was done in September 2023, with seven hives sampled from each district, resulting in a total of fourteen honey samples. The honey was harvested using the pot-puncture technique that is outlined in Mduda et al. (2023d). Subsequently, all honey samples were filtered using a clean food-grade filter cloth, then transferred into amber plastic containers, and stored at 4°C pending laboratory analyses.

### 2.2 Test microorganisms

All test microorganisms used in this study were of standard reference strains from the American Type Culture Collection (ATCC, US). The microbes comprised three Gram-positive bacteria; Methicillin Resistant *Staphylococcus aureus* (MRSA) ATCC 33592, *Staphylococcus aureus* ATCC 6538P and *Bacillus subtilis* ATCC 6633, and two Gram-negative bacteria; *Escherichia coli* ATCC 11229 and *Salmonella typhimurium* ATCC 14028.

### 2.3 Chemicals

Nutrient broth and Mueller Hinton agar were supplied from Himedia Laboratories Private Limited (India). Folin–Ciocalteu phenol reagent, gallic acid (99%), quercetin (98%), aluminium chloride, sodium nitrite, sodium chloride, barium chloride, sodium hydroxide, hydrogen peroxide and methanol were supplied from Glentham Life Sciences (England). Catalase (C100) was supplied from Sigma Aldrich (Germany).

### 2.4 Instrumentation

A class II biosafety cabinet (BSC-1300IIA2-X, BIOBASE) was used to provide controlled environment microbial handling and other sanitary procedures. An incubator (LFZ-TSI-200D, LABFREEZ Instruments) and a UV/Vis spectrophotometer (Cary 60 UV–Vis Spectrophotometer, Agilent Technologies) were also used in this study.

### 2.5 Assessment of the antimicrobial activity of honey

#### 2.5.1 Preparation of inoculum and culture media

Preparation of inoculum and culture media employed the methods outlined in Mduda et al. (2023e). Mueller Hinton agar medium (38 g of Mueller Hinton agar in 1000 mL of distilled water) was prepared and sterilized in an autoclave at 121°C for 15 minutes. The resulting suspension was poured into sterile petri dishes and allowed to solidify at room temperature. Meanwhile, the test microorganisms were inoculated into nutrient broth media (8 g of nutrient broth in 1000 mL of distilled water) in test tubes and then incubated at 37°C for 24 hours. A 0.5 McFarland standard solution was prepared by mixing 0.5 mL of 1.175% (w/v) barium chloride with 99.5 mL of 1% v/v sulfuric acid, and then distributed into screw-capped test tubes. Subsequently, 100 microliters of the inoculated microbe sample from the nutrient broth medium was added to 5 mL of saline, and the concentration was adjusted to 1.5 × 10^8^ colony-forming units (CFU) per milliliter by comparison with the prepared McFarland standard.

#### 2.5.2 Agar-well diffusion assay

The agar-well diffusion assay was carried out according to the procedures outlined in Ewnetu et al. (2013). The bacterial strains were inoculated by streaking the surface of an agar plate with a sterile swab until complete coverage of the agar surface was achieved. Wells were created on the agar plates using a sterile cork borer (6 mm). For the determination of total antibacterial activity, 100µL of 50% (w/v) honey sample in deionized distilled water was added into the agar wells. For non-peroxide activity, 100µL of 50% (w/v) honey sample in catalase solution (10 mg/mL) was used instead (Zainol et al. 2013). The culture plates were then incubated at 37°C for 24 hours. The diameter of inhibition zone was determined by measuring the clear area surrounding the agar wells. Measurements were conducted in both horizontal and vertical directions using a Vernier caliper and recorded in millimeters. Deionized distilled water and Ciprofloxacin (10 µg) were used as negative and positive controls, respectively.

#### 2.5.3 Catalase effectiveness test

Confirmation of catalase removal in honey samples was done following the methods outlined in Zainol et al. (2013). Two honey samples were selected for the test against *S. aureus* ATCC 6538P. Six tubes of test solution were prepared and labeled as follows: tube 1 (50% (w/v) honey solution, 45 mmol/L hydrogen peroxide, and 10 mg/mL catalase solution); tube 2 (50% (w/v) honey solution and 10 mg/mL catalase solution); tube 3 (45 mmol/L hydrogen peroxide and 10 mg/mL catalase solution); tube 4 (50% (w/v) honey solution and 45 mmol/L hydrogen peroxide; tube 5 (50% (w/v) honey solution); and tube 6 (45 mmol/L hydrogen peroxide). These solutions were then tested in the same manner as the agar well diffusion assays on the same plate.

### 2.6 Determination of phytochemical content in honey

#### 2.6.1 Total phenolic content

Determination of phenolic content in honey was conducted following the Folin-Ciocalteau method described by Singleton et al. (1999). Initially, three grams of honey sample were mixed with 30 mL of methanol and subjected to sonication for 15 minutes. Subsequently, the mixture was centrifuged at 9000 rpm, after which the supernatant was carefully decanted and stored at 20°C. Following this step, 2.5 mL of diluted Folin-Ciocalteau phenol reagent and 2 mL of 7.5% sodium carbonate solution were added to 0.5 mL of the extracted sample in a separate tube. The mixture was thoroughly mixed and allowed to stand at room temperature for 2 hours. Absorbance was measured at 765 nm against the blank using a UV–Vis spectrophotometer. A standard calibration curve using gallic acid (0.02 to 0.20 mg/mL) was generated, and the total phenolic content (TPC) was expressed as milligrams of gallic acid equivalent per 100 g of honey (mg GAE/100 g).

#### 2.6.2 Total flavonoid content

Determination of total flavonoid content in honey followed the methods of Zhishen et al. (1999). One mL of the sample solution (5 g of honey in 20 mL of distilled water) was mixed with 4 mL of distilled water and 0.3 mL of 5% sodium nitrite. After a five-minute interval, 0.3 mL of 10% aluminum chloride was introduced into the mixture and allowed to stand for 1 minute. Following this, 2 mL of 1 M sodium hydroxide was added, followed by 2.4 mL of distilled water. The absorbance of the resulting mixture was measured against the blank at 510 nm using a UV/Vis spectrophotometer. A standard calibration curve (0.01– 0.10 mg/mL) was established using quercetin, and the total flavonoid content (TFC) was expressed as milligrams of quercetin equivalent per 100 g of honey (mg QE/100 g).

### 2.7 Data analysis

One-way analysis of variance (ANOVA) and a Tukey post-hoc test were employed for the comparison of the total and non-peroxide antibacterial activities of honey samples from the two locations. Non-metric multidimensional scaling (NMDS) ordination plot was created using Euclidean similarity index to highlight the similarities in the total and non-peroxide antibacterial activities. Data were log transformed before generating the NMDS plot to fit them in the same scale. ANOVA was also used to compare susceptibility of the bacterial strains to the studied honey samples. Two sample t-test was conducted to compare the phytochemical content of honey samples from the two locations. Pearson’s correlation coefficient was performed to evaluate association between diameters of inhibition zones and phytochemical content of honey. Data analysis was done using PAleontological STatistics (PAST) Software V. 4.03 and graphs were plotted by GraphPad Prism V. 9.5.1.

## 3.0 Results and Discussion

### 3.1 Total versus non-peroxide antibacterial activity of stingless bee honey

Results of the total antibacterial activity of stingless bee honey against the test microbes are presented in Table 1. Honey samples exhibited antibacterial activity against all bacterial strains except for two samples from Kibiti (KB02 and KB03) which failed to inhibit the growth of *E. coli* ATCC 11229. The highest total antibacterial activity was observed in sample KB05 from Kibiti against *S. aureus* ATCC 6538P (Table 1).

**Table 1.** Diameters of inhibition zones (mm) produced by honey samples before treatment with catalase against five bacterial strains.

| Test microbes | MRSA<br>ATCC 33592 | *S. aureus*<br>ATCC 6538P | *B. subtilis*<br>ATCC 6633 | *E. coli* ATCC 11229 | *S. typhimurium*<br>ATCC 14028 |
| --- | --- | --- | --- | --- | --- |
| | Mean ± SD | Mean ± SD | Mean ± SD | Mean ± SD | Mean ± SD |
| SH01 | 13.0 ± 1.0 | 16.0 ± 0.0 | 14.9 ± 0.3 | 11.7 ± 0.6 | 11.3 ± 1.2 |
| SH02 | 15.0 ± 0.5 | 15.7 ± 0.6 | 12.5 ± 1.1 | 11.0 ± 1.0 | 13.3 ± 1.2 |
| SH03 | 16.0 ± 0.5 | 14.3 ± 0.6 | 12.3 ± 0.6 | 10.5 ± 0.3 | 12.0 ± 0.0 |
| SH04 | 13.0 ± 1.0 | 13.7 ± 1.2 | 14.0 ± 0.3 | 10.5 ± 0.5 | 11.7 ± 0.6 |
| SH05 | 14.0 ± 1.0 | 14.3 ± 0.6 | 13.8 ± 0.3 | 12.5 ± 0.5 | 13.0 ± 0.0 |
| SH06 | 13.5 ± 0.5 | 13.5 ± 1.2 | 12.5 ± 0.5 | 11.0 ± 0.6 | 16.0 ± 1.5 |
| SH07 | 13.5 ± 0.3 | 14.8 ± 0.3 | 14.4 ± 0.6 | 12.0 ± 0.0 | 13.3 ± 1.5 |
| KB01 | 12.0 ± 0.0 | 13.3 ± 1.5 | 13.0 ± 1.1 | 10.0 ± 0.0 | 11.7 ± 0.6 |
| KB02 | 11.0 ± 0.0 | 14.2 ± 0.6 | 11.2 ± 0.3 | *6.0 ± 0.0 | 12.3 ± 0.6 |
| KB03 | 12.5 ± 1.5 | 13.3 ± 0.6 | 11.0 ± -.3 | *6.0 ± 0.0 | 10.5 ± 0.5 |
| KB04 | 13.5 ± 0.5 | 13.7 ± 1.2 | 12.5 ± 0.6 | 14.5 ± 0.5 | 12.8 ± 0.3 |
| KB05 | 11.8 ± 0.3 | 16.3 ± 0.6 | 12.3 ± 0.6 | 11.0 ± 0.0 | 13.3 ± 0.6 |
| KB06 | 12.0 ± 0.0 | 13.7 ± 0.6 | 13.5 ± 0.3 | 10.0 ± 0.0 | 12.0 ± 1.0 |
| KB07 | 12.5 ± 0.5 | 13.7 ± 0.6 | 12.3 ± 0.6 | 10.2 ± 0.8 | 12.0 ± 0.0 |

Samples with codes SH and KB originated from Siha and Kibiti districts, respectively. Size of the agar well is 6.00 mm. Values with \* sign indicate no microbial inhibition.

Honey exhibits potent antimicrobial activity due to various attributes such as its low pH, high osmolarity and the presence of hydrogen peroxide and non-peroxide components (Mandal and Mandal 2011). Previous studies have reported pH of *M. ferruginea* honey ranging between 3.8 and 4.9 (Mokaya et al 2022, Mduda et al. 2023a) which is low enough to be inhibitory to bacteria. Honey acidity is influenced by the presence of organic acids, particularly gluconic acid which is the dominant acid in honey (Dardón et al. 2013). However, pH is raised when honey is diluted making it less effective as an antimicrobial factor (Mduda et al. 2013e). Additionally, *M. ferruginea* honey has lower sugar content (70.3 – 73.9 °Brix) and higher water content (26.1 - 28.8%) compared to *A. mellifera* honey (Mokaya et al. 2022, Mduda et al. 2023a), resulting to low osmolarity.

Honey generates hydrogen peroxide via the enzyme glucose oxidase which oxidizes glucose to gluconic acid and hydrogen peroxide (Mandal and Mandal 2011). Antimicrobial potency due to hydrogen peroxide (peroxide activity) is the most common in many types of honey, with maximum activity when the honey is diluted (Almasaudi 2021). The downside of the peroxide activity is that hydrogen peroxide can be easily destroyed by heat or in the presence of catalase enzyme (Mduda et al. 2023e). In that regard, the effectiveness of hydrogen peroxide as an antimicrobial agent is limited when honey is mixed in bodily fluids (Almasaudi 2021, Mduda et al. 2024).

Non-peroxide antibacterial activity was assayed after treating the studied honey samples with catalase enzyme and the results are presented in Table 2. Catalase effectiveness test showed that the enzyme was effective in removing all hydrogen peroxide molecules from the honey samples and its activity was not affected by other components (Supplementary table). The highest non-peroxide antibacterial activity was observed in sample SH05 against *S. aureus* ATCC 6538P. Similar to the results of total antibacterial activity, two honey samples from Kibiti failed to inhibit the growth of *E. coli* ATCC 11229. No significant difference was observed between the total and non-peroxide antibacterial activity against all bacterial strains for honey samples from Siha and Kibiti (Fig. 1). Honey samples from both locations retained the majority of antibacterial activity after treatment with catalase (Fig. 2, Table 3). Similarly, the NMDS plot (Fig. 3) shows minimal differences between the total and non-peroxide antibacterial activity based on the extent to which the convex hulls overlap.

**Fig. 1.**
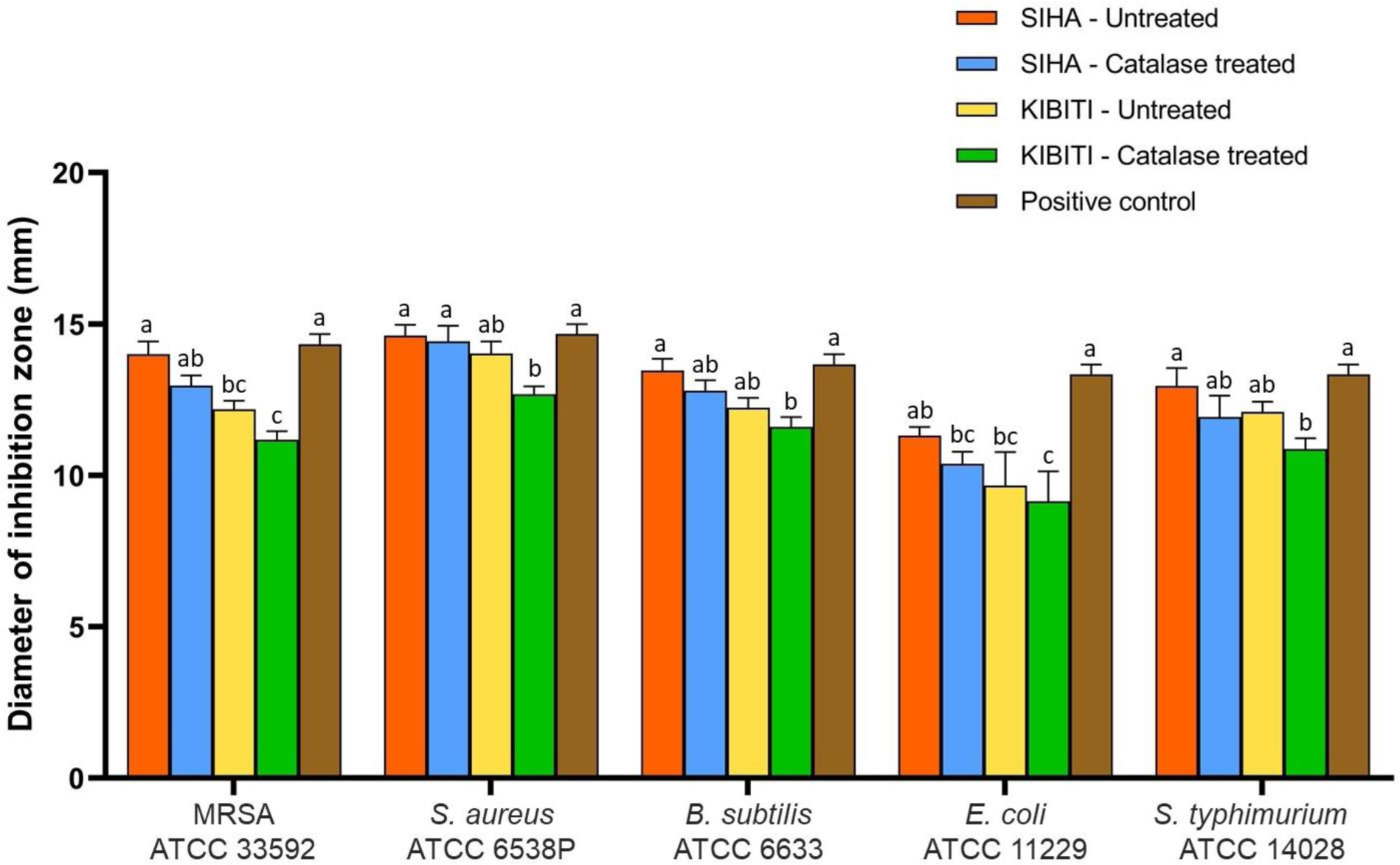
Grouped bar-graphs showing mean diameters of inhibition zones of untreated (total activity) and catalase-treated (non-peroxide activity) honey samples against the bacterial strains. Ciprofloxacin (10 µg) was used as a positive control. Superscripts with different letters within the same group indicate significant differences in diameters of inhibition zones.

**Fig. 2.**
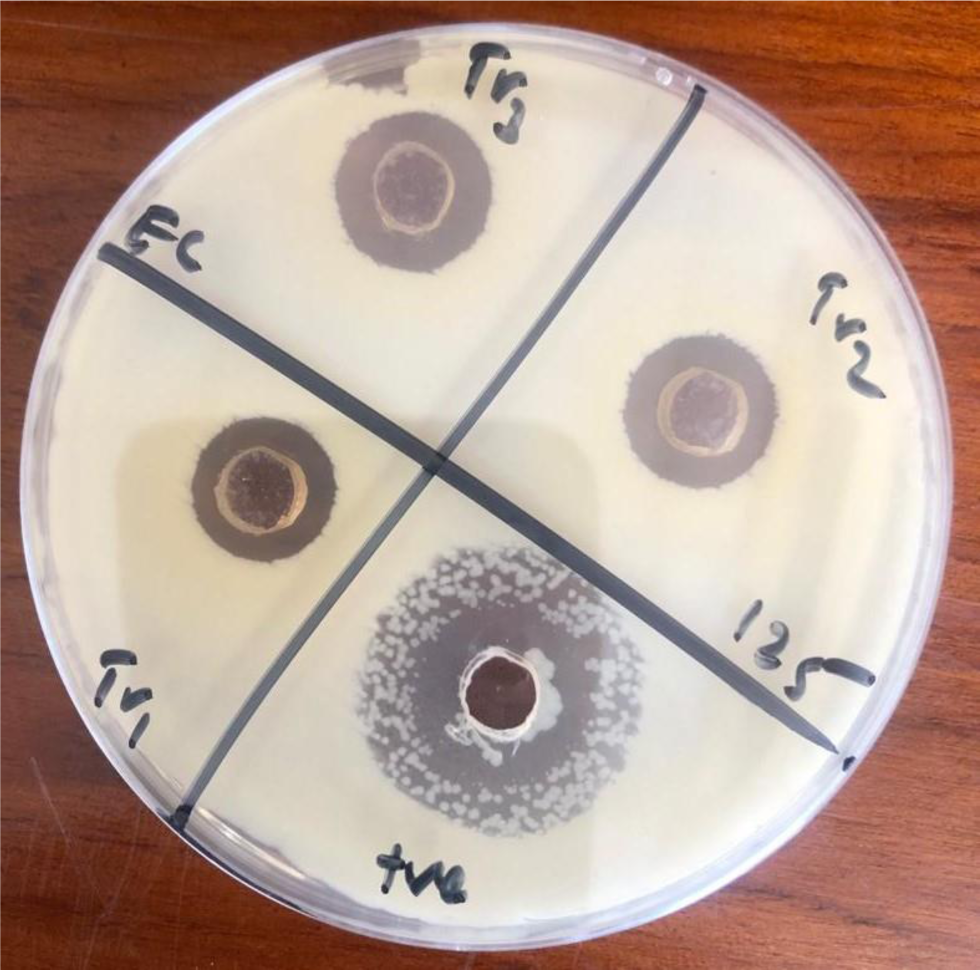
A representative agar plate showing zones of inhibition produced by stingless bee honey. Wells in Tr1 and Tr2 received catalase-treated honey (Non-peroxide activity) while Tr3 received untreated honey (Total activity).

**Fig. 3.**
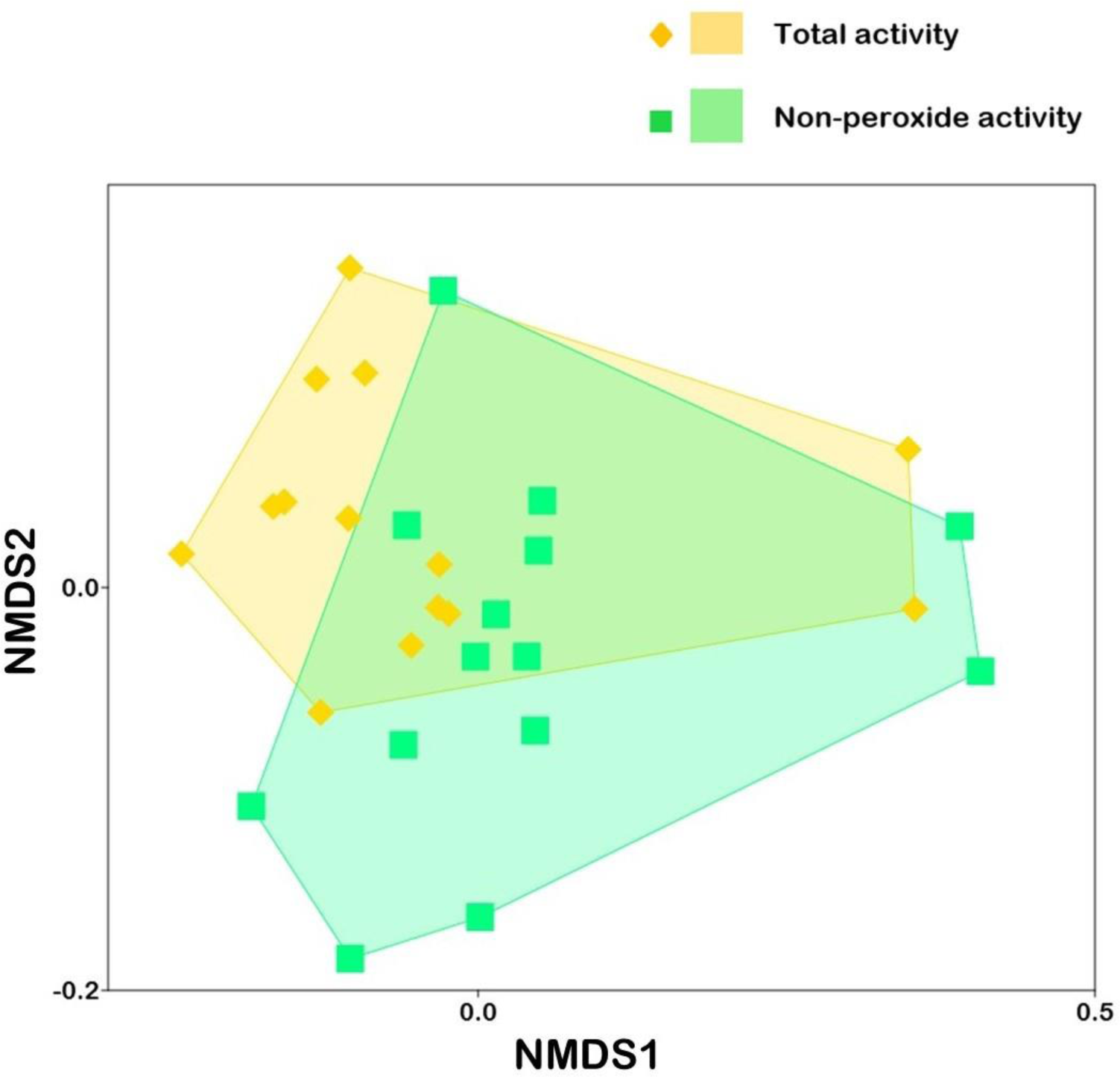
Non-metric multidimensional scaling (NMDS) ordination plot showing honey treatments exhibiting total and non-peroxide antibacterial activity. Overlapping of convex hulls indicate the degree of similarity between sample treatments. Stress value of the NMDS plot is 0.159.

**Table 2.** Diameters of inhibition zones (mm) produced by catalase-treated honey samples against five bacterial strains.

| Test microbes | MRSA<br>ATCC 33592 | *S. aureus*<br>ATCC 6538P | *B. subtilis*<br>ATCC 6633 | *E. coli* ATCC<br>11229 | *S. typhimurium*<br>ATCC 14028 |
| --- | --- | --- | --- | --- | --- |
| | Mean ± SD | Mean ± SD | Mean ± SD | Mean ± SD | Mean ± SD |
| SH01 | 12.3 ± 0.3 | 16.0 ± 0.0 | 14.2 ± 0.3 | 10.3 ± 0.6 | 9.8 ± 0.3 |
| SH02 | 13.8 ± 0.8 | 15.0 ± 0.5 | 12.3 ± 0.6 | 10.3 ± 1.2 | 12.5 ± 0.5 |
| SH03 | 14.5 ± 0.3 | 14.0 ± 1.0 | 11.7 ± 1.1 | 8.7 ± 0.6 | 11.8 ± 0.3 |
| SH04 | 12.0 ± 0.0 | 13.0 ± 1.0 | 13.3 ± 0.6 | 9.7 ± 0.6 | 11.0 ± 0.0 |
| SH05 | 13.0 ± 0.5 | 16.0 ± 0.0 | 13.3 ± 0.6 | 12.0 ± 1.0 | 12.5 ± 0.5 |
| SH06 | 12.8 ± 0.8 | 12.5 ± 0.5 | 11.8 ± 1.1 | 10.3 ± 0.6 | 15.5 ± 0.5 |
| SH07 | 12.5 ± 0.5 | 14.5 ± 0.5 | 13.0 ± 0.6 | 11.3 ± 0.6 | 10.5 ± 0.8 |
| KB01 | 10.5 ± 0.5 | 12.5 ± 0.5 | 12.3 ± 0.6 | 9.7 ± 0.6 | 10.5 ± 0.5 |
| KB02 | 10.0 ± 0.0 | 13.0 ± 1.0 | 10.7 ± 1.1 | *6.0 ± 0.0 | 11.0 ± 1.0 |
| KB03 | 11.0 ± 0.5 | 12.8 ± 0.3 | 10.3 ± 0.6 | *6.0 ± 0.0 | 9.3 ± 0.6 |
| KB04 | 11.8 ± 0.3 | 13.0 ± 1.0 | 11.7 ± 0.6 | 13.7 ± 0.6 | 10.5 ± 0.5 |
| KB05 | 11.5 ± 0.5 | 11.5 ± 0.5 | 11.7 ± 0.3 | 9.7 ± 0.6 | 12.5 ± 0.5 |
| KB06 | 11.3 ± 0.3 | 13.8 ± 0.3 | 12.8 ± 0.3 | 9.0 ± 1.0 | 11.3 ± 0.3 |
| KB07 | 12.3 ± 0.8 | 12.3 ± 0.8 | 11.7 ± 0.6 | 10.0 ± 0.0 | 11.0 ± 0.0 |

Samples with codes SH and KB originated from Siha and Kibiti districts, respectively. Size of the agar well is 6.00 mm. Values with \* sign indicate no microbial inhibition.

**Table 3:**
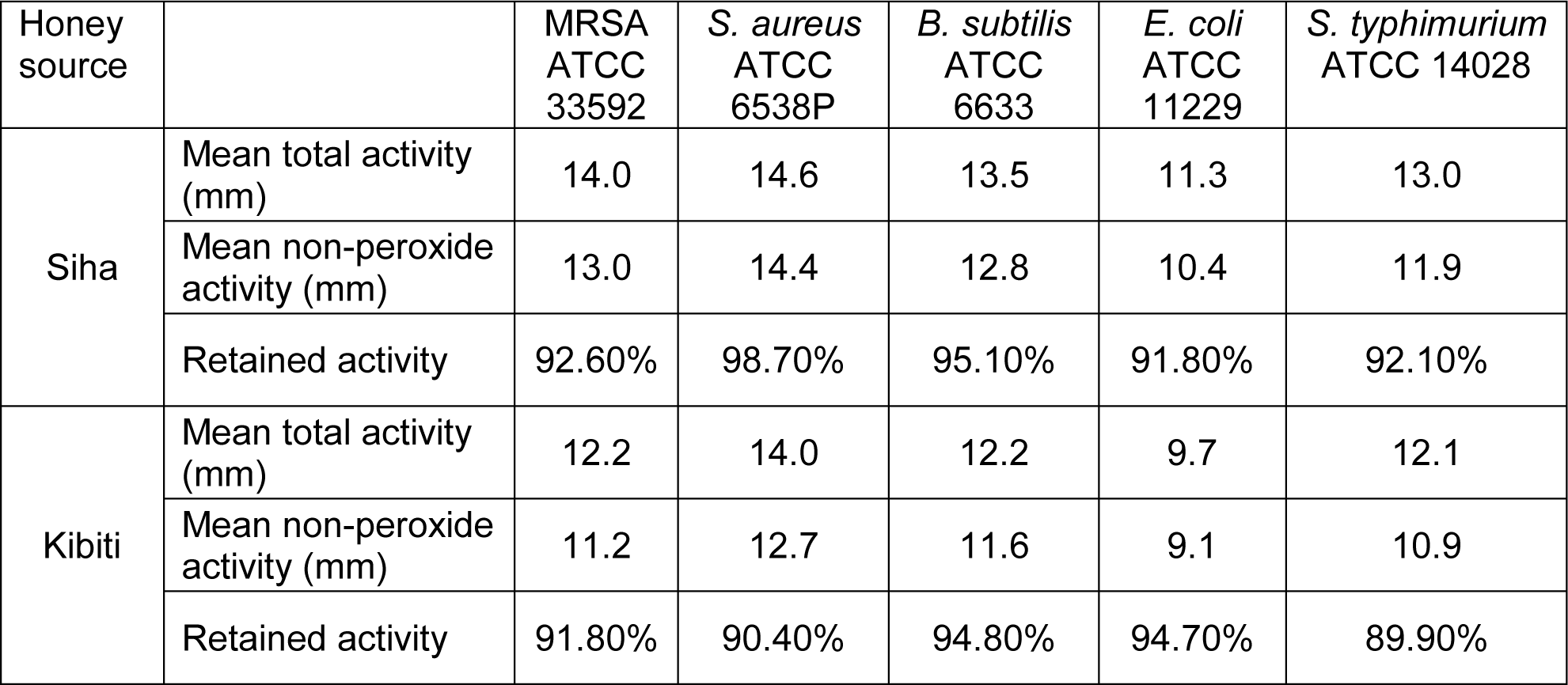
Antibacterial activity retained after treatment of honey samples with catalase enzyme.

Previous studies have highlighted the prevalence of non-peroxide antibacterial activity in stingless bee honey (Temaru et al. 2007, Zainol et al. 2013, Jibril et al. 2020). Honey samples from 14 stingless bee species displayed remarkable non-peroxide antibacterial activity against gram positive and gram-negative bacterial strains (Temaru et al. 2007). Further, Jibril et al. (2020) reported that stingless bee honey retained 98.9% of the antibacterial activity after treatment with catalase (Jibril et al. 2020). Contrary, non-peroxide activity is uncommon in *Apis mellifera* honey except for special honey types such as Manuka honey (Johnston et al. 2018). For example, honey samples from various origins in UK and Denmark had no detectable non-peroxide activity despite exhibiting broad-spectrum total antibacterial activity (Sulaiman et al. 2012, Matzen et al. 2018). Additionally, the removal of hydrogen peroxide by catalase resulted in substantial decrease in the antibacterial activity of *A. mellifera* honeys from Western Australia (Roshan et al. 2017).

Several non-peroxide components have been reported to contribute to the antibacterial activity of honey. The most extensively studied are methylglyoxal and methyl syringate which are found predominantly in Manuka honey (Johnston et al. 2018, El-Senduny et al. 2021, Hossain et al. 2023). Honey also contains a diverse array of phytochemicals including polyphenols which are derived from the nectar of flowers (Mduda et al. 2023c). Polyphenols such as phenolic acids and flavonoids can exhibit antibacterial potency by interfering with the bacterial cell functioning, disrupting cell growth and effecting cell lysis (Shehu et al. 2016, Kumar Singh et al. 2019). Additionally, honey may contain bee derived proteins such as the antimicrobial peptides which may also contribute to its antimicrobial potency (Almasaudi 2021).

### 3.2 Phytochemical content of stingless bee honey

Stingless bee honey exhibited remarkable levels of total phenolic (197.0 – 263.1 mg GAE/100 g) and flavonoid content (118.5 – 156.7 mg QE/100 g) (Fig. 4). Polyphenols can play a critical role in the non-peroxide antibacterial activity of honey (Tuksitha et al. 2018). Previous studies reported that antimicrobial activity of honey strongly correlated with both TPC and TFC (Sousa et al. 2016, Mduda et al. 2023e). In contrast, it was observed in this study that both TPC and TFC lacked significant correlation with the diameters of inhibition zones against any of the bacterial strains (Fig. 5). When comparing the two locations, honey samples from Kibiti produced smaller mean diameters of inhibition zones (Fig. 1), despite having significantly higher amounts of total phenolic and flavonoid content. Tuksitha et al. (2018) also observed that honey samples with the highest TPC failed to inhibit the growth of gram negative bacteria. In addition, honey samples from Brazil and Scotland showed no correlation between TPC and antimicrobial activity against a variety of bacterial strains including *Shigella dysentery*, *Salmonella typhimurium*, *Staphylococcus aureus* and *Bacillus cereus* (Bueno-Costa et al. 2016, Fyfe et al. 2017). These results indicate that the antimicrobial activity displayed by the studied honey samples may be attributed to specific polyphenolic compounds present in honey rather than the total content. The amount and types of polyphenols present in honey vary depending on the botanical source, geographical location as well as the bee species origin (Mduda et al. 2023e). Specific phenolic acids such as Syringic acid, Chlorogenic acid, Caffeic acid and Hydroxycinnamic acids, and flavonoids such as Naringenin, Apigenin, Quercetin and Myricetin, have been identified to exhibit broad spectrum antibacterial activity (Jibril et al. 2019). Hence, future research should focus on identifying the specific polyphenols and other bioactive compounds responsible for the antimicrobial potency observed in Tanzanian stingless bee honeys.

**Fig. 4.**
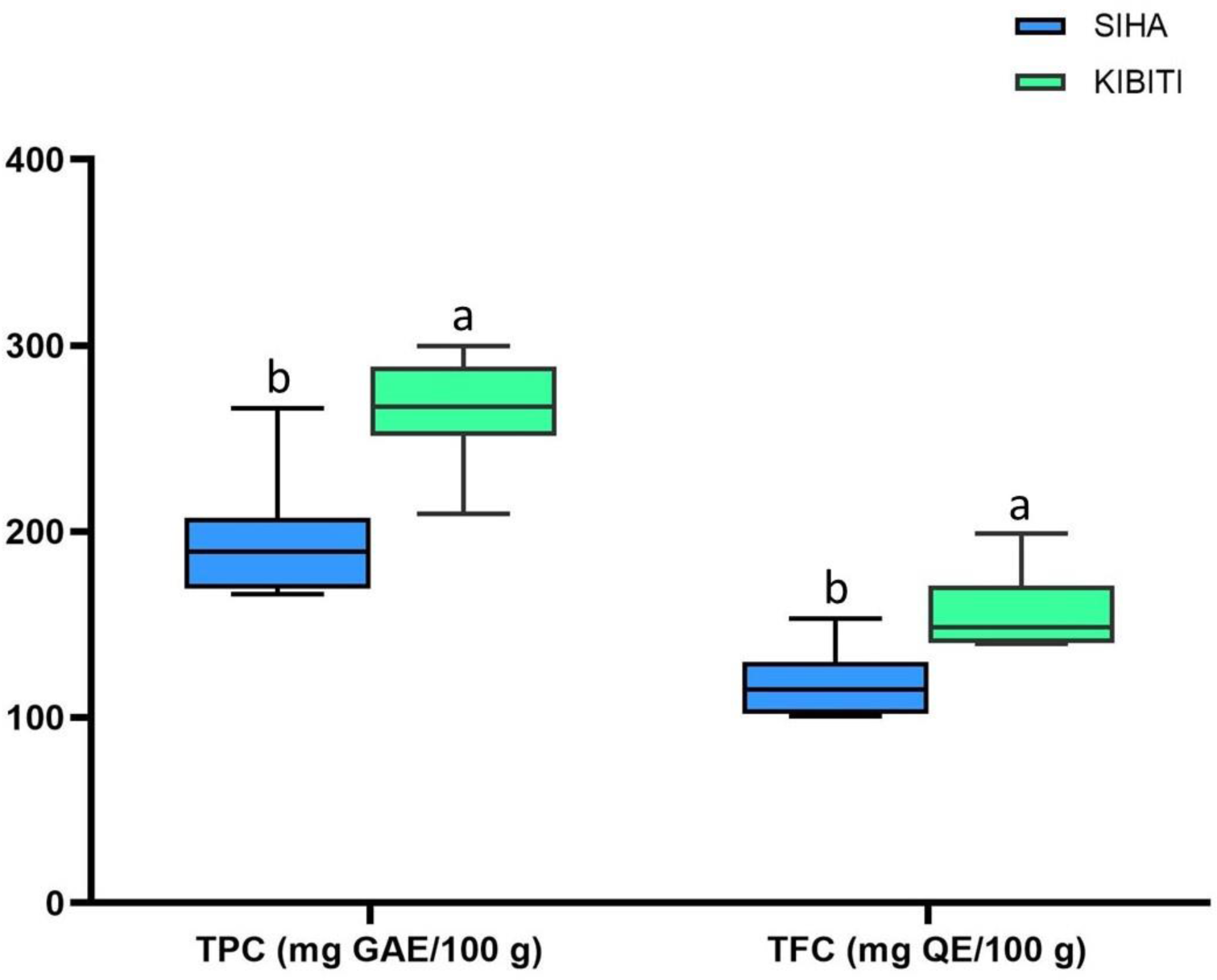
Grouped bar-graph showing total phenolic and flavonoid content of stingless bee honey from the studied locations.

**Fig. 5.**
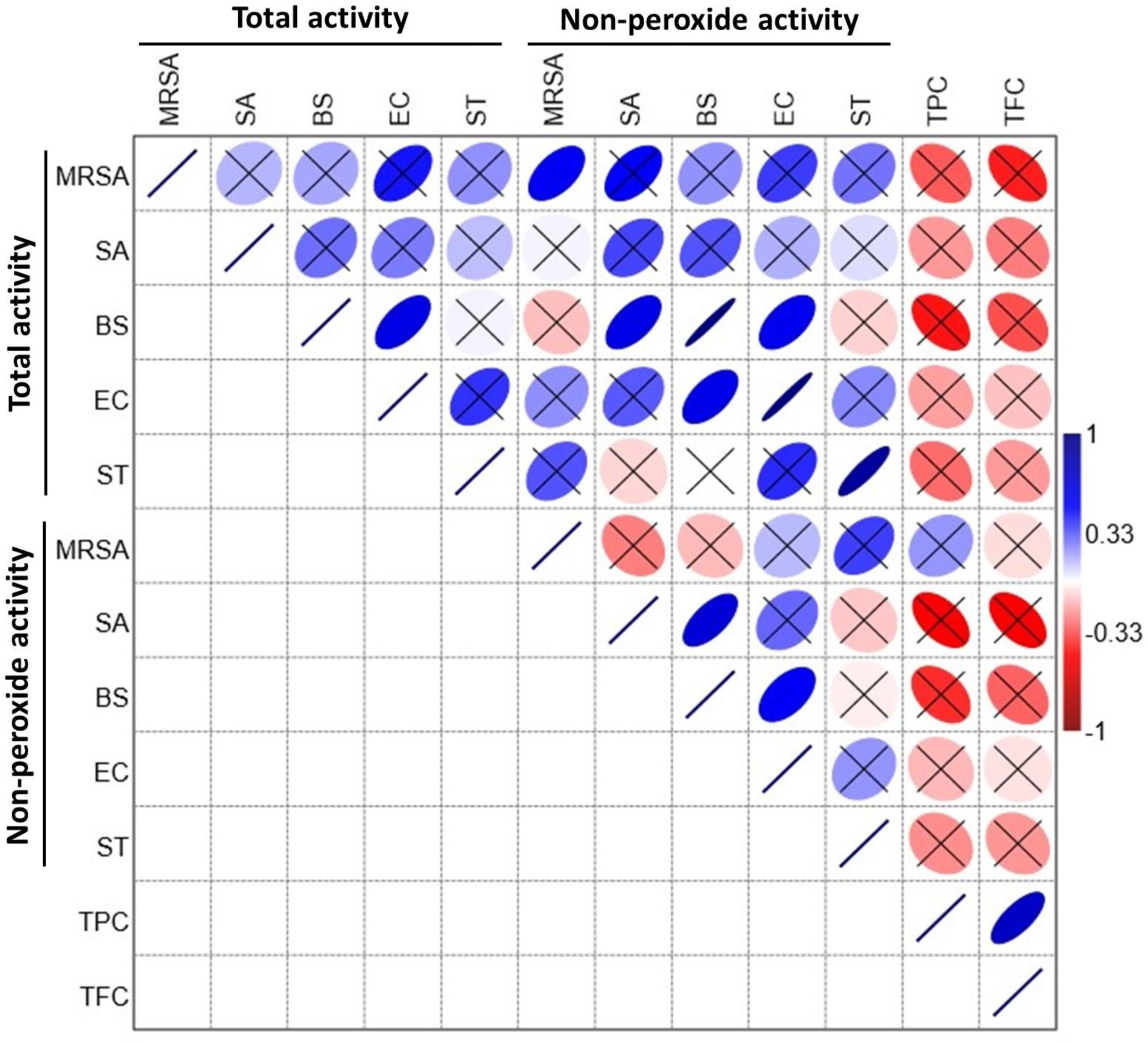
Correlation matrix showing pairwise Pearson’s coefficients among variables. Crossed boxes (×) indicate correlations which are not statistically significant (p > 0.05). MRSA = Methicillin resistant *S. aureus* ATCC 33592, SA *= S. aureus* ATCC 6538P, BS = *B. subtilis* ATCC 6633, EC = *E. coli* ATCC 11229, ST = *S. typhimurium* ATCC 14028, TPC total phenolic content, TFC = total flavonoid content.

### 3.3 Microbial susceptibility to stingless bee honey

The bacterial strains displayed different levels of susceptibility to both untreated and catalase treated honey samples (Fig. 4). Stingless bee honey was more effective in inhibiting the growth of gram-positive bacteria compared to gram-negative bacteria. The largest and smallest diameters of inhibition zones were observed against *S. aureus* ATCC 6538P and *E. coli* ATCC 11229, respectively. Several authors have also reported stingless bee honey to be more effective against gram-positive bacteria compared to gram-negative bacteria (Zainol et al. 2013, Domingos et al. 2021, Mduda et al. 2023e). Malaysian stingless bee honey displayed remarkably high total and non-peroxide activity against *S. aureus* than the standard medicinal Manuka honey (Zainol et al. 2023). Similarly, honey samples from two *Scaptotrigona* species effectively inhibited the growth of two MRSA strains, while being least effective against strains of *E. coli* (Nishio et al. 2016). In contrast, Ng. et al (2020) reported Malaysian stingless bee honey to be highly effective against *E. coli*. Variation in microbial susceptibility to honey may result from differences in growth rate and cell-wall permeability to antimicrial components (Dżugan et al. 2020). Additionally, honey samples from different origins may comprise bioactive compounds with different effects on bacterial cells.

**Fig. 6.**
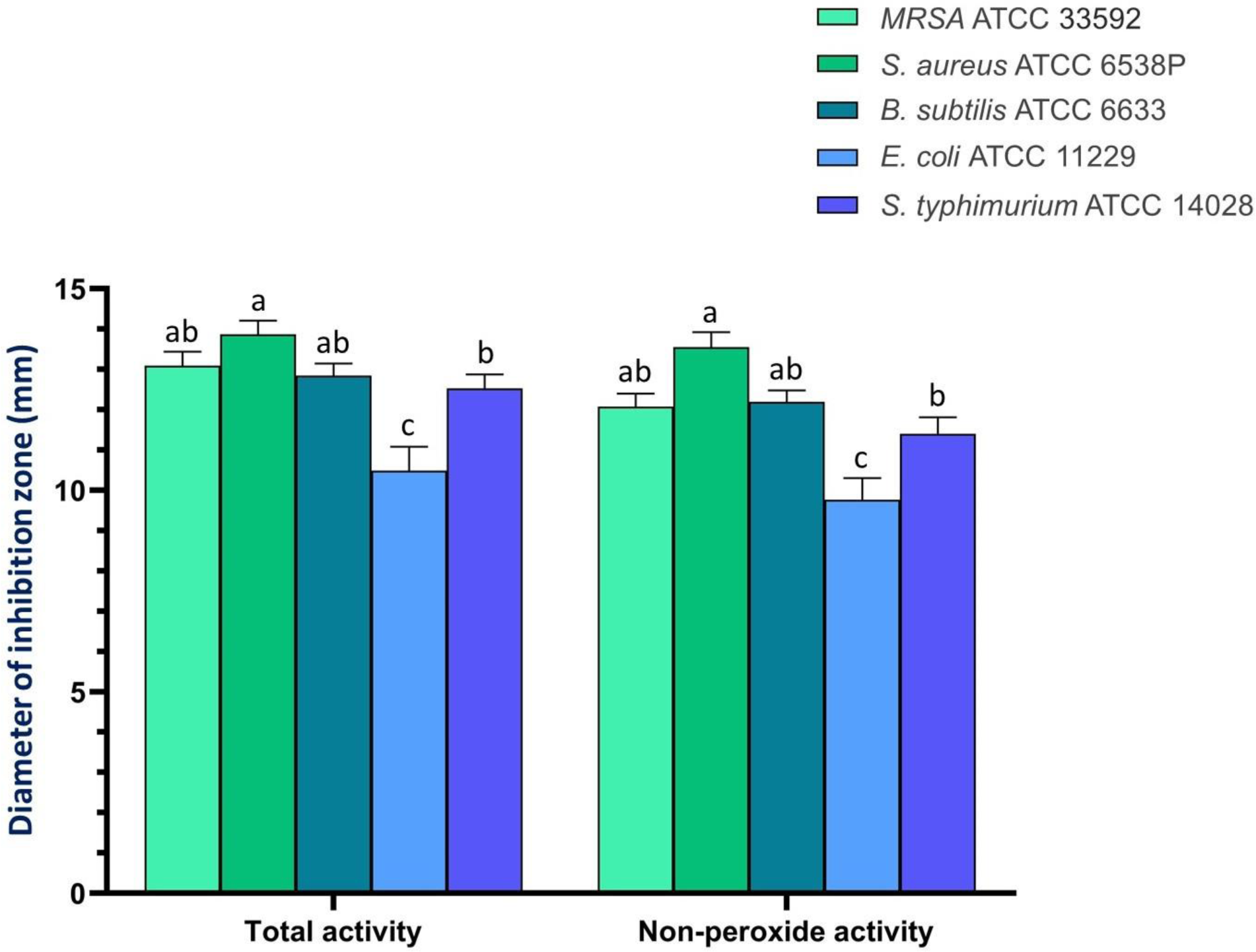
Grouped bar-graph showing differences in microbial susceptibility to the untreated (total activity) and catalase-treated (non-peroxide activity) honey samples.

## 4.0 Conclusion

The studied honey samples demonstrated broad-spectrum antibacterial activity against common pathogens. Primarily, the antibacterial efficacy of *M. ferruginea* stemmed from its non-peroxide component, indicating substantial therapeutic value. Its notable effectiveness against Methicillin Resistant *S. aureus* advocates it as a potent natural solution for the treatment of wound infections and drug-resistant pathogens. Further investigations are needed to elucidate the mechanisms and bioactive compounds underlying the observed activity in order to ascertain its clinical potential for treating bacterial infections including resistant strains.

## Acknowledgements

The authors would like to thank the beekeepers who provided honey samples for this study.

## Data availability statement

All data supporting the findings of this study are available within the article and its supplementary files.

