## Supplementary Table for "Non-peroxide antibacterial activity of *Meliponula* (*Axestotrigona*) *ferruginea* honey from Tanzania"

**Supplementary table**: Results of the catalase effective test showing diameter of inhibition zones (mm) of treatments against *Staphylococcus aureus* ATCC 6538P

| Treatments | Without catalase | | With catalase | |
|---|---|---|---|---|
| | SH01 | KB02 | SH01 | KB02 |
| Honey solution (50% w/v) + Hydrogen peroxide (45 mmol/L) | 26.5 ± 0.5 | 25.8 ± 0.3 | 16.0 ± 0.5 | 13.5 ± 0.3 |
| Honey solution (50% w/v) | 16.3 ± 0.3 | 14.3 ± 0.3 | 15.8 ± 0.8 | 13.3 ± 0.3 |
| Hydrogen peroxide (45 mmol/L) | 29.3 ± 3 | | *6.0 ± 0.0 | |

Values are recorded in mean ± SD. Size of the agar well is 6.00 mm. Values with * sign indicate no microbial inhibition.
